# Enhanced COVID-19 data for improved prediction of survival

**DOI:** 10.1101/2020.07.08.193144

**Authors:** Wenhuan Zeng, Anupam Gautam, Daniel H Huson

## Abstract

The current COVID-19 pandemic, caused by the rapid world-wide spread of the SARS-CoV-2 virus, is having severe consequences for human health and the world economy. The virus effects individuals quite differently, with many infected patients showing only mild symptoms, and others showing critical illness. To lessen the impact of the pandemic, one important question is which factors predict the death of a patient? Here, we construct an enhanced COVID-19 dataset by processing two existing databases (from Kaggle and WHO) and using natural language processing methods to enhance the data by adding local weather conditions and research sentiment.

**Author summary:** In this study, we contribute an enhanced COVID-19 dataset, which contains 183 samples and 43 features. Application of Extreme Gradient Boosting (XGBoost) on the enhanced dataset achieves 95% accuracy in predicting patients survival, with country-wise research sentiment, and then age and local weather, showing the most importance. All data and source code are available at http://ab.inf.uni-tuebingen.de/publications/papers/COVID-19.

## Introduction

The current COVID-19 pandemic, caused by the rapid world-wide spread of the SARS-CoV-2 virus, is affecting many aspects of society, in particular human health, but also social issues [1, 2], mental health and the economy [3]. Medical researchers, and researchers from different scientific fields, including immunology, genetics and bioinformatics, are studying the pandemic to find ways to slow its progression. Machine learning approaches are being utilized to understand aspects of the problem.

To date, most machine learning research on COVID-19 has used supervised learning methods or deep learning [4, 5] to investigate which might be the important features to predict a predefined outcome. Running such approaches on the publicly available datasets is associated with difficulties that are due to the fact that features are collected depending on the needs of the data provider, which can be a source of bias. In particular, features that have high predictive value for the outcome for an infected patient, might be missing. Generally speaking, the presence or absence of features will impact the accuracy of a model.

The currently available COVID-19 data is missing features and we explore the effect of this by adding a number of features that might be important, so as to determine how this affects the accuracy of the model.

We used data on patients that tested positive for the virus and added new features based on (1) how different countries responded to the pandemic in terms of research sentiment (so as to calculate a weighted average polarity score for research abstracts per country) and (2) the local weather conditions when the patient was probably infected. We found that age is one of the most important factors when we have not incorporated these additional features based on the initial data.

However, after the addition of two new features, country-specific research sentiment, followed by local weather and age, came out to be the most important features. Recent publications suggest that the weather, as represented by the variables temperature and humidity, plays a role in COVID-19 [6] and SARS [7].

To summarize, our main contributions are as follows:

- We demonstrate how to construct an enhanced set of COVID-19 features using additional available information.
- Using this enhanced dataset, we show that the Extreme Gradient Boosting (XGBoost) method achieves 95% accuracy in predicting a patient’s survival.
- We show that country-specific research sentiment, followed by age and local weather and are the most important features.

## Materials

We first compiled an initial dataset by combining data from two sources.

### Kaggle Novel Corona Virus 2019 dataset

We used data from Kaggle available at its Novel Corona Virus 2019 Dataset portal (https://www.kaggle.com/sudalairajkumar/novel-corona-virus-2019-dataset), which contains day-level information on COVID-19 cases. The dataset includes features such as ID, age, sex, city, province and country. The dataset file (COVID19_open_line_list.csv) was downloaded on 30/03/2020. All the rows that do not contain a value in the outcome column were dropped, resulting in 183 patient data rows out of 13,174. The final dataset contains 183 patients from 16 countries. Further processing was carried out on this dataset (S1 File).

### WHO COVID-19 database

We downloaded a database of literature on COVID-19 from the World Health Organization (WHO) web site (https://www.who.int/emergencies/diseases/novel-coronavirus-2019/global-research-on-novel-coronavirus-2019-ncov), on April 13, 2020. Of the 5,354 downloaded entries, we kept only those whose “Journal Name” and “DOI” fields were not blank, which resulted in 4,683 publications in 590 journals. We then analyzed these publications to determine the authors’ institute and country (S2 File).

### COVID-19 enhanced dataset

In this paper, we present an enhanced COVID-19 dataset, which is based on the above described initial database. The data is enhanced by adding features that reflect the local weather and research sentiment in the country of the infected person, as described in the following (S3 File).

### Addition feature construction

Database construction was performed as outlined in Fig 1. It has been demonstrated that there is a link between environmental factors and the development of COVID-19 [8]. Indeed, it seems reasonable to suspect that the weather conditions play a role.

**Fig 1.**
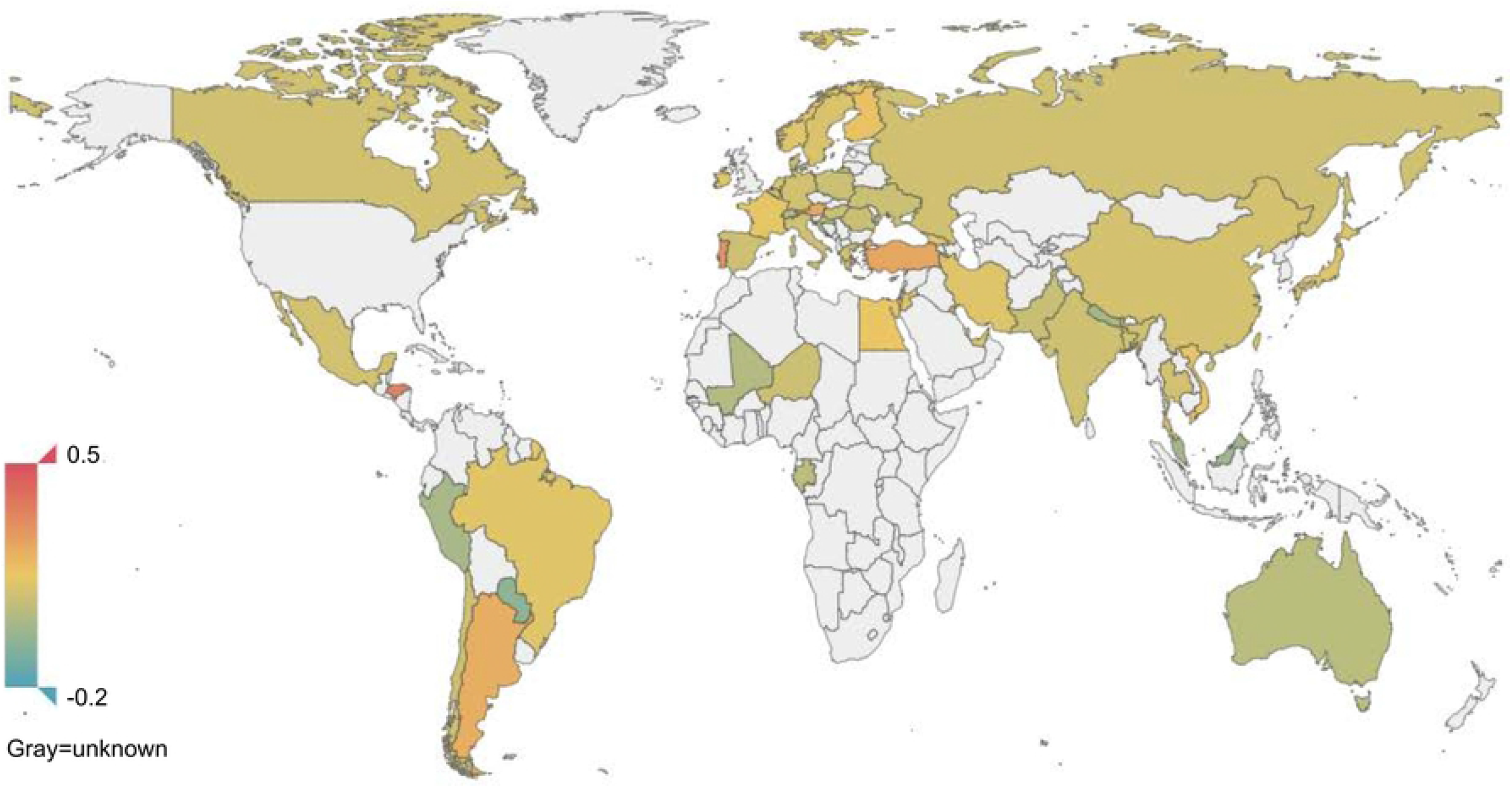
Construction of database. Enhancement. The Kaggle COVID-19 database is filtered for patients for which the outcome has been recorded, and then, for these items, the weather is determined using the https://www.wunderground.com website. The WHO COVID-19 literature database is filtered for items for which both a journal name and DOI are provided, and these are post-processed so as to obtain a country-wise research sentiment polarity score. XGBoost is then trained and run on both the initial and the enhanced data and the accuracy of survival prediction is shown to be 85% and 95%, respectively.

Hence, we collected temperature, humidity and a text description of the weather for the city where the patient lives from the Weather Underground website (https://www.wunderground.com/). To take into account that the incubation period of the virus is approximately 14 days, we collected this data for a date 14 days before the patient exhibited relevant symptoms (as recorded in the Novel Corona Virus 2019 dataset).

For a given country, we assume that the researchers’ attitude toward COVID-19 will reflect the response capacity of the country, to some extent. For journal publications obtained from the WHO database, we extracted the author’s institution with the help of the paper’s DOI. We then applied sentiment analysis on each abstract so as to obtain a polarity score for every abstract, and we then calculated an weighted average polarity score for each country. This feature was added to the enhanced dataset. Fig 2 displays the weighted average polarity score inferred for different countries.

**Fig 2.**
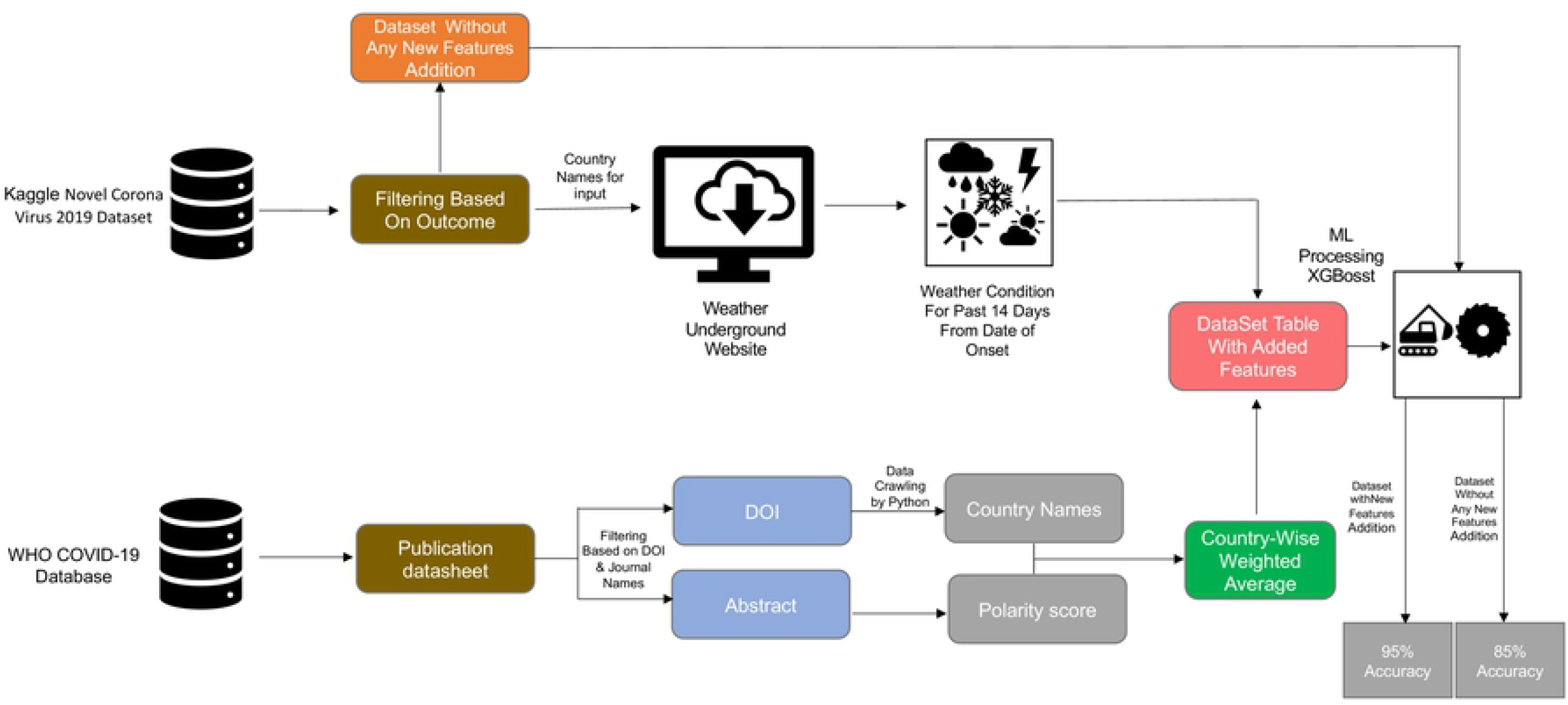
Sentiment polarity of abstract. Average research sentiment polarity score of research, for different countries. Based on a sentiment analysis of abstracts of papers published on COVID-19.

### Data processing

The features obtained from the Kaggle Novel Corona Virus 2019 dataset include both categorical variables and discrete variables. Each item in the dataset contains the variables sex, age, the time interval between the patient’s onset date, confirmed infected date and admission date, symptoms description, infection reason and outcome. We will refer to this as the *initial* dataset.

We added local weather variables (temperature, humidity, climate description) and the weighted polarity score of country’s research attitude. We will refer to the result of this as the *enhanced* dataset.

To prepare for analysis with XGBoost (as discussed below), we tokenized all multi-value text features, such as symptom description or climate description, into three-dimensional embedding vectors, used label encoding on categorical variables such as infected reason, as shown in Table 1.

**Table 1.** Example of multi-value text features.

| Features | Example |
| --- | --- |
| symptom description | fever, cough, sputum |
| weather description | Fair, Light Rain Shower, Cloudy |

We assigned the constant −999 to all missing values. After filtering for samples that have a valid outcome value, we obtain 183 samples.

### Data statistics

Due to the relatively small size of the two datasets, we split each dataset into a training set and a testing set in the proportion of 8:2. As shown in Table 2, the original dataset is typically imbalanced, and after applying the Synthetic Minority Over-sampling Technique (SMOTE) [9] on the minority group, the ratio of the positive samples and the negative sample is 1:5. Note that, for technical reasons, here positive samples refer to patients that died.

**Table 2.** Sampling statistics for the enhanced dataset.

| Statistic | Enhanced data | After oversampling |
| --- | --- | --- |
| Positive samples | 15 | 33 |
| Negative samples | 168 | 168 |
| Total | 183 | 201 |

## Methods and experiment

### Sentiment analysis

A number of papers have studied the forecasting of pandemics using natural language processing on data obtained from various social media [10–12]. Along these lines, we performed sentiment analysis on the abstracts of research papers (associated with COVID-19) using the Python package Textblob (https://github.com/sloria/TextBlob), which operates by analyzing text content and assigns emotional values to word based on matches to a built-in dictionary.

### Machine learning algorithm

Our aim is to predict whether the patient will survive the infection, based on either the initial dataset or the enhanced dataset.

We use the Extreme Gradient Boosting (XGBoost) [13] method to address this. XGBoost is a powerful member of the gradient boosting family, which is designed to perform well on sparse features, and is known to perform well on Kaggle tasks, This approach avoids overfitting using its built-in L1 and L2 regularization on the target function:

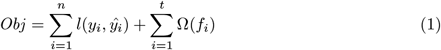

As an additive model, XGBoost consists of *k* base models, and in most cases we choose the tree model as its base model. Suppose, for the *k*-th of *t* iterations, that we train the tree model f_k_(x), then

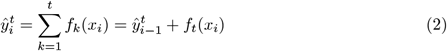

is the estimate result of the sample *i* after *t* times’ iteration. During construct of each tree, XGBoost minimizes the objective function with regularization term introduced in Eq (1) in the split phase of each node. In each tree, we calculate the *Gain* of the feature and choose the tree who has the biggest value as the leaf node to be split:

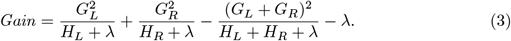

### Implementation

In this study, we ran the XGBoost algorithm on two different datasets, namely the initial dataset and the enhanced dataset, the latter containing additional features representing local weather and research sentiment, as illustrated in Fig 1.

To obtain the model with the best capacity for prediction, we used a grid search for model tuning. Each subtree in our model is a simple tree whose maximum depth is 3. The learning rate was 0.01. During the training step, we randomly sampled the columns of each tree according to a ratio of 0.5.

### Results

We evaluated the algorithm’s performance by calculating each model’s classification accuracy. The accuracy of the model created by using the initial dataset (no added features) is 85%, whereas using the enhanced dataset (with added features), the model’s accuracy is 95%.

The method we chose to evaluate the importance score of feature is based on counting the number of times that a feature occurred in a tree. The feature importance for both datasets is shown in Fig 3. For the initial dataset, age plays a more important role than other features. For the model based on the enhanced dataset, weighted average research sentiment polarity score is more important than age, whereas the level of importance of weather is similar to that of age.

**Fig 3.**
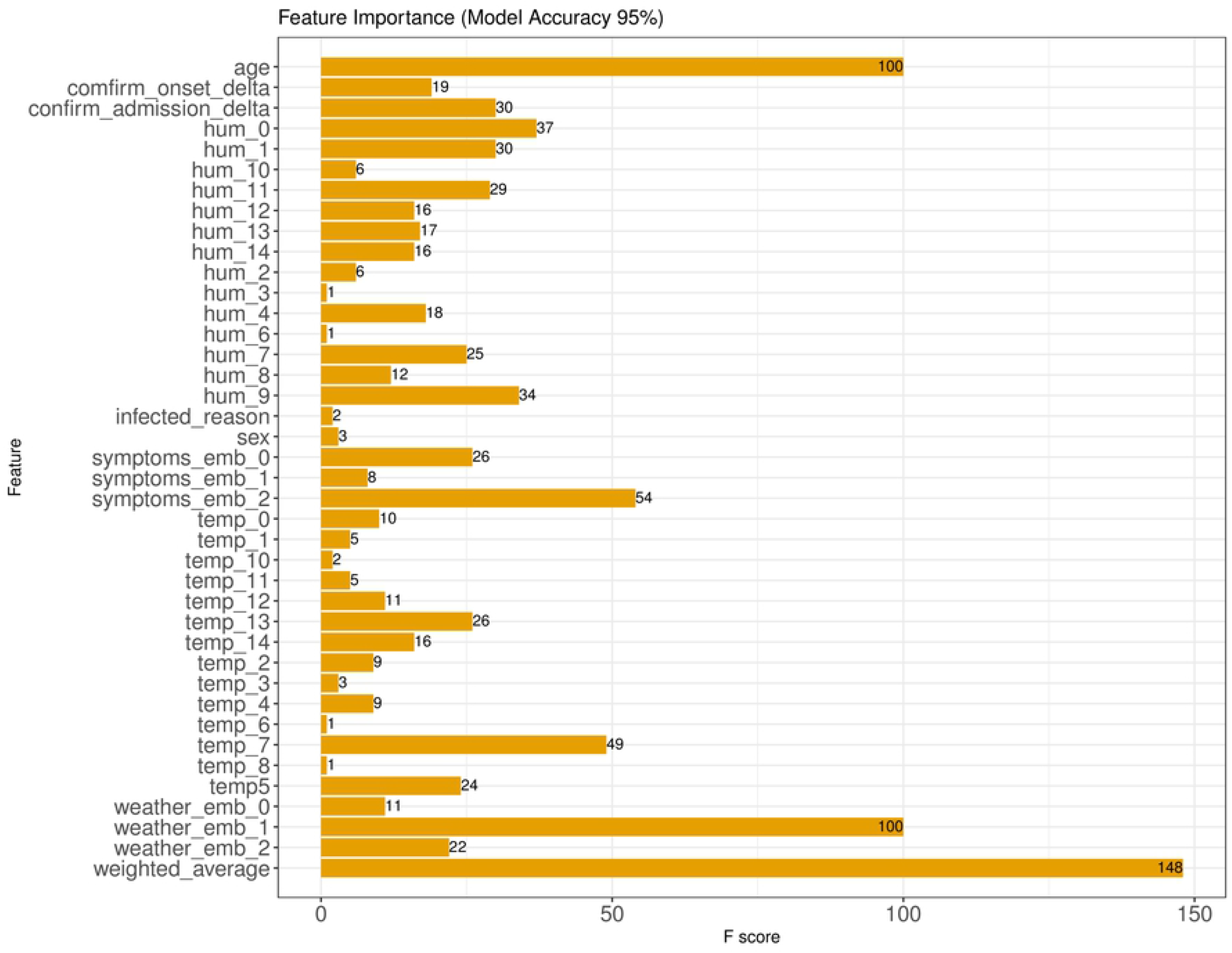
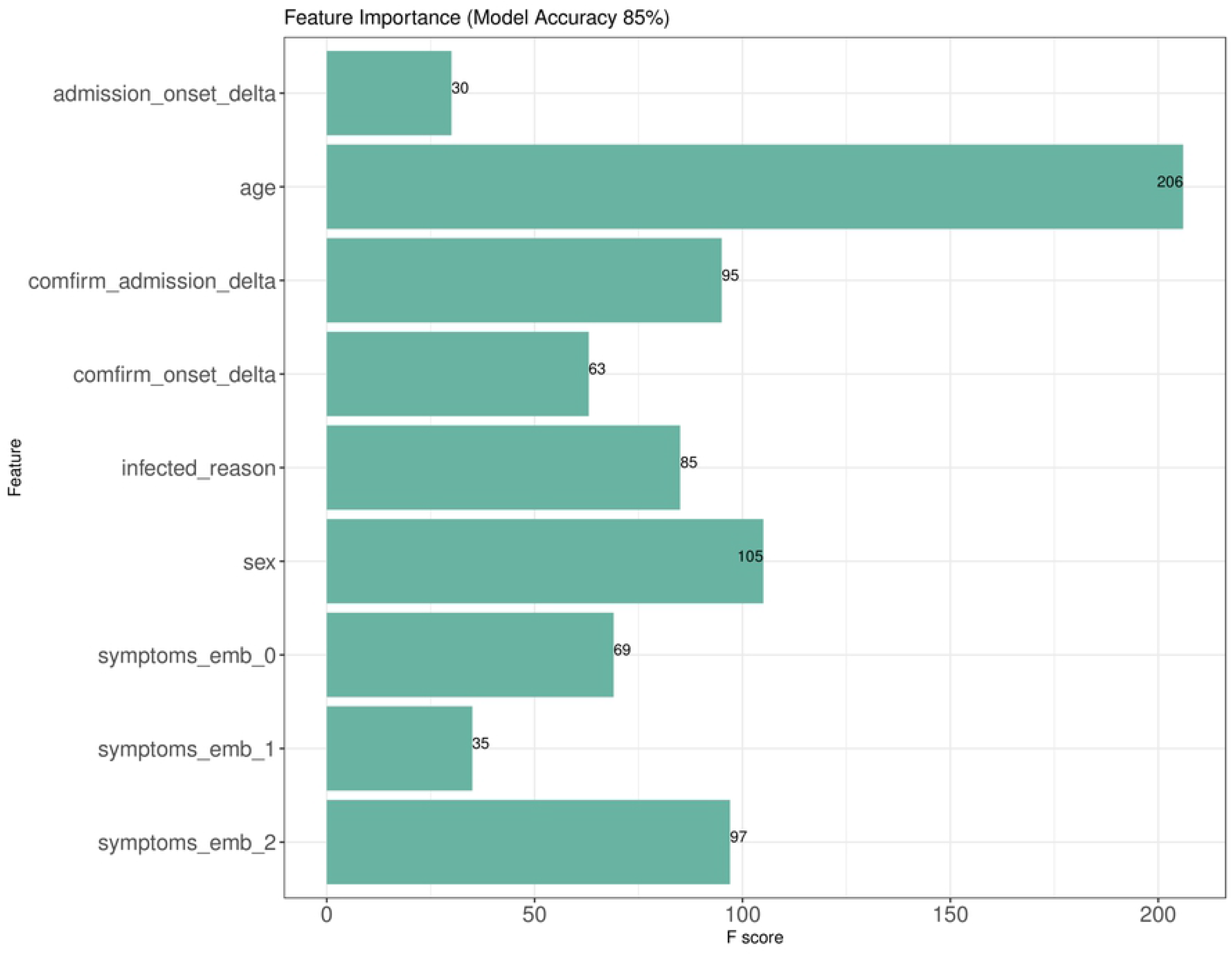
Feature scores on initial and enhanced datasets. (A) Feature scores on the initial dataset. (B) Feature scores on the enhanced dataset.

## Conclusion

The performance of machine learning methods depends on the amount and quality of available features. For our analysis, we can say that the current publicly available data is poor. First, the data is quite sparse and there are too few features. Here we see that by enhancing the dataset, the accuracy of survival prediction can be increased by 10%.

Our study shows how one might enhance a dataset by adding informative features if they are not available in the original dataset. Here we demonstrated this for country-wise research sentiment and local weather. Local weather conditions has been implicated as an important feature in the existing research.

Our analysis confirms the observation that age is an important factor for survival of COVID-19. However, in the data considered here, the total number of deaths above age 60 were 8, and 16 survived or were still alive, while in the age group between 40-60 there were 2 deaths and 36 alive or survived. Hence, linking mortality to a particular age group is not be appropriate based on the current result. While this analysis suggests that elderly have a higher risk of death, which has already been observed [14, 15], saying mortality is associated with old age is probably generally true for any infectious disease. Age is one of the confounding factors that could be responsible for enhanced COVID-19 mortality rate, so more emphasis should be be taken for the elderly care [16, 17].

For the model based on the enhanced dataset, the weighted average of research sentiment, followed by weather and age, appear as the most important features, and account for the increase in the accuracy of the model. This confirms that environmental conditions play a role. Also, it suggests the research sentiment might reflect a countries ability to tackle the disease.

Finally, this analysis suggests that enhancing a dataset, rather than just analyzing the originally given features, might lead to a better prediction of the particular outcome.

## Supporting information

**S1 File. Initial COVID-19 dataset 183 cases.**

**S2 File. Processed WHO publication data.**

**S3 File. Enhanced COVID-19 dataset.**

